# Low-intensity transcranial focused ultrasound suppresses pain by modulating pain processing brain circuits

**DOI:** 10.1101/2022.12.07.519518

**Authors:** Min Gon Kim, Kai Yu, Chih-Yu Yeh, Raghda Fouda, Donovan Argueta, Stacy Kiven, Yunruo Ni, Xiaodan Niu, Qiyang Chen, Kang Kim, Kalpna Gupta, Bin He

**Affiliations:** Department of Biomedical Engineering, Carnegie Mellon University, Pittsburgh, PA 15213; Department of Medicine, University of California at Irvine, Irvine, CA 92697; Department of Medicine, University of Pittsburgh, Pittsburgh, PA 15261; Department of Bioengineering, University of Pittsburgh, Pittsburgh, PA 15261; Neuroscience Institute, Carnegie Mellon University, Pittsburgh, PA 15213

**Keywords:** Chronic pain, Sickle cell disease, Behaviors, Electrophysiology, Non-invasive brain stimulation, Transcranial focused ultrasound

## Abstract

There is an urgent and unmet clinical need to develop non-pharmacological interventions for chronic pain management due to the critical side effects of opioids. Low-intensity transcranial focused ultrasound is an emerging non-invasive neuromodulation technology with high spatial specificity and deep brain penetration. Here, we developed a tightly-focused 128-element ultrasound transducer to specifically target small mouse brains, employing dynamic focus steering. We demonstrate that transcranial focused ultrasound stimulation at pain processing brain circuits can significantly alter pain-associated behaviors in mouse models in vivo. Our findings indicate that a single-session focused ultrasound stimulation to the primary somatosensory cortex (S1) significantly attenuates heat pain sensitivity in wild-type mice and modulates heat and mechanical hyperalgesia in a humanized mouse model of chronic pain in sickle cell disease. Results further revealed a sustained behavioral change associated with heat hypersensitivity by targeting deeper cortical structures (e.g., insula) and multi-session focused ultrasound stimulation to S1 and insula. Analyses of brain electrical rhythms through electroencephalography demonstrated a significant change in noxious heat hypersensitive- and chronic hyperalgesia-associated neural signals following focused ultrasound treatment. Validation of efficacy was carried out through control experiments, tuning ultrasound parameters, adjusting inter-experiment intervals, and investigating effects on age, gender, genotype, and in a head-fixed awake model. Importantly, transcranial focused ultrasound was shown to be safe, causing no adverse effects on motor function and brain neuropathology. In conclusion, the rich experimental evidence validates the ability of novel focused ultrasound neuromodulation to suppress pain, presenting significant translational potential for next-generation chronic pain treatment without adverse effects.

**Key points:**

- Novel non-invasive neuromodulation of brain’s pain processing circuits with submillimeter spatial precision for pain management
- Transcranial focused ultrasound significantly modulates pain-related behaviors and brain electrical rhythms of pain in humanized SCD mice

## Introduction

The multidimensional impact of chronic pain has closely correlated with various aspects of the quality of life and an increased risk of developing physical and mental health problems^1,2^. While opioids remain the mainstay for treating chronic pain associated with underlying conditions such as cancer or sickle cell disease (SCD), the higher death rates due to drug overdose have been recognized as a significant health crisis^3^. Consequently, prioritizing the management of chronic pain with non-pharmacologic treatments becomes crucial to mitigate the risks associated with opioids, including drug abuse, overdose, and addiction^3,4^. Several non-pharmacologic neuromodulation methods have been explored^5–7^; however, the clinical application of these methods is constrained by invasive surgical procedures and/or challenges in stimulating specific and deeper brain structures^8–9^.

Low-intensity transcranial focused ultrasound (tFUS) is an emerging neuromodulation technology with unparalleled spatial specificity and penetration depth. Capable of non-invasively modulating neuronal activities in specific brain regions^10–15^, tFUS holds promise in treating neurological disorders^16–18^. Notably, tFUS has been applied to managing pain by targeting specific brain circuits: stimulating the right anterior thalamus for antinociception in healthy individuals^19^, targeting the posterior frontal cortex to induce mood improvement in chronic pain patients^20^, modulating thalamus and periaqueductal gray (PAG) regions to reduce heat pain response in non-human primates^21^, and stimulating the PAG to attenuate formalin-mediated nociceptive activity in the spinal cord dorsal horn in rats^22^. Despite these promising applications in pain research, several fundamental questions remain. Firstly, it is unknown whether low-intensity tFUS stimulation, applied to certain pain processing brain circuits, can effectively alter pain-related behaviors. This is crucial considering that noxious stimuli activate multiple brain areas^23,24^, including the primary and secondary somatosensory cortex (S1 and S2), insula, thalamus, anterior cingulate cortex, and PAG. Secondly, the brain region specificity of the tFUS analgesic effect needs further investigation of neural oscillations for quantification and validation. Thirdly, establishing effective and safe pain management remains elusive due to lack of translatable evidence and limited utilization of pain assays in human interventional experiments^25,26^. Alternatively, well-established rodent models of chronic pain in SCD namely BERK and Townes exhibit similarity to human genetic and pathological disease expressing over 99% human sickle hemoglobin^27,28^ and offering potential for translational pain research^29,30^. Consequently, an urgent need arises to address the aforementioned critical aspects for an effective and safe approach to pain treatment in the humanized mouse model.

In this study, we present a transformative approach employing low-intensity tFUS neuromodulation for chronic pain management. Our investigation involved stimulating specific pain processing brain circuits in both wild-type mice and humanized SCD mice which exhibit constitutive chronic hyperalgesia similar to that observed in sickle cell human patients. We hypothesize that tFUS stimulation can change (i.e., deteriorate or ameliorate) pain-associated behaviors triggered by various noxious stimuli through modulating the activities of specific brain circuits involved in pain processing (Fig. 1). Leveraging a custom-designed, small footprint, 128-element highly-focused random array ultrasound transducer, we achieved a high spatial specificity and focus steering capability for ultrasound stimulation in the small mouse brain. By harnessing this array transducer, we systematically studied the short-term and sustained effects of single- and multi-session tFUS stimulation with varying parameters and targeting specific brain regions. Our investigation delved into how tFUS stimulation can alter pain-associated behaviors evoked by noxious heat, cold, and mechanical stimuli. Utilizing novel ultra-flexible scalp electroencephalography (EEG) recordings, we demonstrate how tFUS modulates neural oscillations in the specific pain-processing brain circuit underlying chronic pain and noxious heat stimuli. Importantly, to ascertain the specificity of the modulatory effect to tFUS stimulation, we conducted a comprehensive set of control experiments. Lastly, behavior and histological evaluations were performed for rigorous safety assessment. The combination of these approaches provides a robust foundation for exploring the potential of tFUS as a novel and effective modality for chronic pain management.

**Figure 1.**
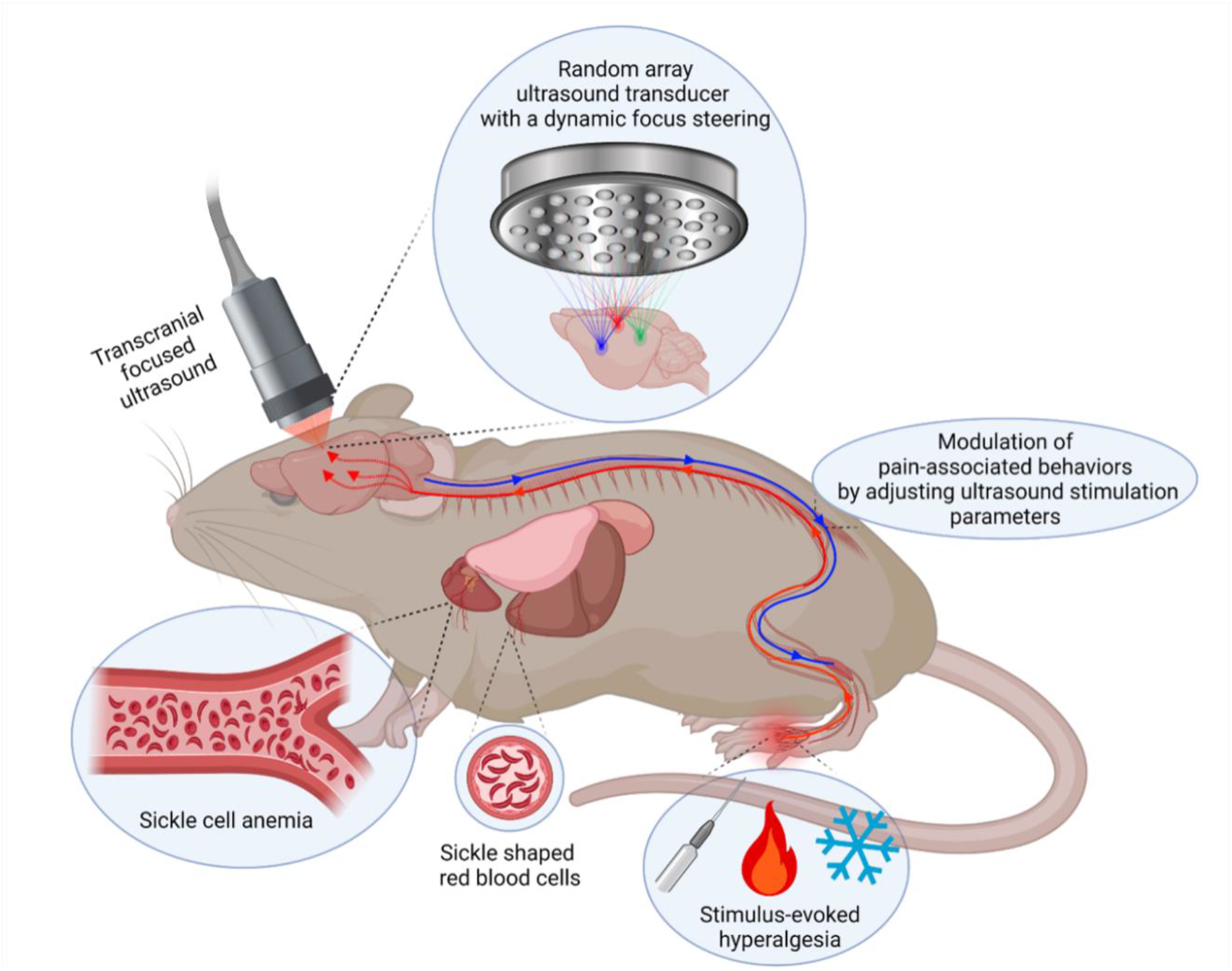
The conceptual diagram illustrates the hypothesis of treating chronic and stimulus-evoked hyperalgesia through transcranial focused ultrasound (tFUS) stimulation in a humanized mouse model of chronic pain in sickle cell disease (HbSS-BERK). The modulation of pain-associated behaviors, such as amelioration or deterioration, is proposed to be achieved by selectively stimulating specific pain processing brain circuits. This is facilitated using a 128-element highly-focused random array ultrasound transducer capable of a dynamic focus steering in HbSS-BERK mice. The diagram captures the idea of targeted neuromodulation to influence pain responses in the context of sickle cell disease.

## Methods

### Animals

Male and female wild-type mice (C57BL/6J), male and female humanized transgenic HbSS-BERK with control mice (bred in house), and female HbSS-Townes mice were used. HbSS-BERK mice demonstrate several features of SCD including chronic pain; and control mice expressing normal human HbA (human α and β globins and no mouse globins)^27,31^ (see Supplementary Information for details). All studies were approved by Institutional Animal Care and Use Committee of Carnegie Mellon University and Long Beach VA medical Centre and complied with National Institutes of Health guidelines.

### Experimental design for behavioral testing and EEG measurement

Fig. 2A illustrates the behavioral experimental setup, incorporating various types of behavioral tests, tFUS stimulation with multiple control groups, and video assessment with subsequent data analysis. In Fig. 2B, a study paradigm for EEG measurements is presented to simultaneously investigate how tFUS stimulation influences intrinsic brain activities and heat stimulus-evoked activities. The tFUS, transmitted by the random array ultrasound transducer (H276), was directed onto the right hindlimb representation area of the S1 (S1HL), followed by noxious heat stimulation (≤ 47 °C) to the plantar surface of the contralateral hind paw. In these behavioral and EEG experiments, controls were implemented, including a negative control group (no tFUS stimulation, maintaining the same experimental procedures) and a sham-treated group (applying tFUS to a control brain structure near the bregma). These controls were essential to rule out potential confounding factors from the observed tFUS-mediated modulatory effects (Fig. 2C). After characterizing the customized H276 probe driven by a Vantage 128 ultrasound system (Supplementary Fig. S1-S3; see Supplementary Information for details), the tFUS beam was electronically steered and applied with controllable parameters, including a tone-burst duration (TBD) of 200 µs, pulse repetition frequency (PRF) of 40 Hz and 3 kHz, ultrasound duration (UD) of 100 ms and 400 ms, inter-sonication interval (ISoI) of 2-second and 4-second, and total sonication time of 10-minute, 20-minute, and 1-hour (Fig. 2D).

**Figure 2.**
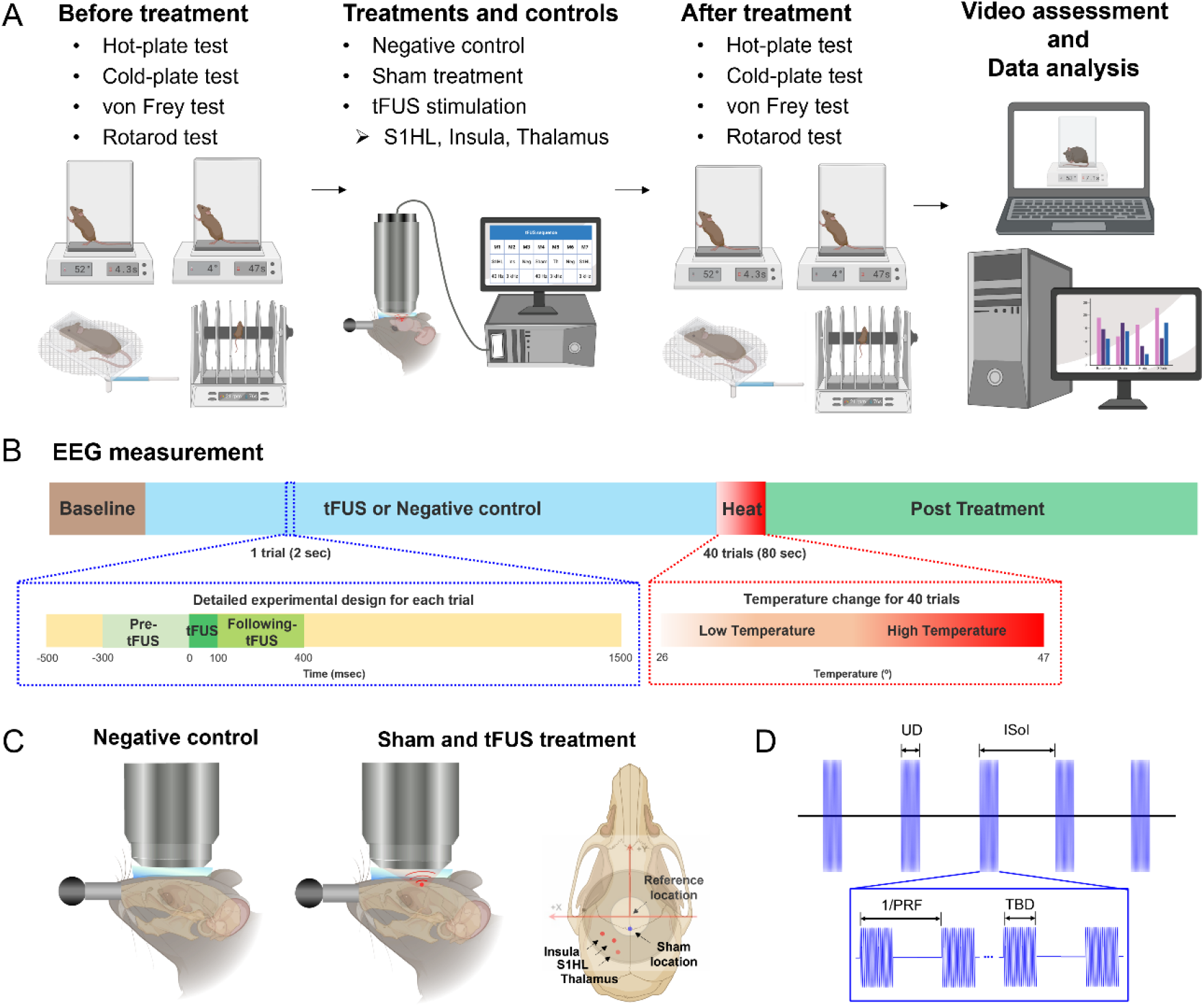
Experimental design. (A) Schematic of behavioral experimental setup: The setup comprises various behavioral tests, tFUS stimulation with multiple control groups, and video assessment with subsequent data analysis. Nocifensive reactions to heat, cold, and mechanical stimuli, as well as motor performance, were recorded and evaluated to investigate the effects of tFUS stimulation at different brain locations, including S1HL, insula, and thalamus. (B) Study design for electroencephalography (EEG) measurements: EEG measurements were designed to examine both intrinsic brain activity and heat stimulus-evoked brain activity along with tFUS stimulation. (C) Control experiments: To determine the specificity of the modulatory effect to tFUS stimulation at a particular brain circuit, the effects of negative control (no tFUS stimulation, maintaining experimental procedures) and sham treatment (applying tFUS to a control brain structure near the targeted brain location) were examined. (D) tFUS parameters: The 128-element random array transducer was customized to steer ultrasound focus. Targeted brain structures were subject to tFUS with specific parameters, including a fundamental frequency of 1.5 MHz, tone-burst duration (TBD) of 200 µs, pulse repetition frequency (PRF) of 40 Hz and 3 kHz, ultrasound duration (UD) of 100 ms and 400 ms, inter-sonication interval (ISoI) of 2 s and 4 s, and total sonication time of 10 min, 20 min and 1 h.

All behavioral experiments were conducted during the light cycle of the animals and recorded on video for further validation (refer to the Supplementary Information for detailed procedures of the pain-associated behavior assessment). For heat hyperalgesia, the temperature of the hot-plate (Ugo Basile, PA, USA) was pre-determined at 52 °C for HbSS-BERK mice (53 °C for wild-type mice) and hind paw withdrawal latency (hPWL) was determined through blind assessments. For cold hyperalgesia, the temperature of the cold-plate (Ugo Basile, PA, USA) was pre-determined at 4 °C based on a prior study^31,32^ and hind paw withdrawal frequency (hPWF) over a 2-minute period was blindly assessed. For mechanical hyperalgesia, minimum threshold of von Frey filaments (Stoelting Co, IL, USA) was calculated using the up-down method^28,33^ and represented as a hind paw withdrawal threshold (hPWT). Motor performance was assessed using a rotating rod (Ugo Basile, PA, USA) programmed to gradually increase revolutions per minute (rpm) from 3 to 72 rpm over 5-minute^34^. The integral of time spent on the rod (i.e., latency to fall) was measured. In the behavioral analysis, the unilateral effect of tFUS was assessed^35^, and behavioral responses to the hot, cold, and mechanical stimuli were calculated using the following metrics: Δ hPWL (s) = contralateral (right) hPWL-ipsilateral (left) hPWL, Δ hPWF = contralateral hPWF-ipsilateral hPWF, and Δ hPWT = contralateral hPWT-ipsilateral hPWT.

### Head-restrained awake naturally walking mice

The head fixation platform (Mobile HomeCage, Neurotar Ltd, Helsinki, Finland) was used to investigate the potential influence of light anesthesia on observed pain-related behaviors in wild-type mice (see Supplementary Information for details).

### Histology

Brain sections, following a multi-session treatment for 14 days with 1-hour per day, underwent staining with hematoxylin and eosin (H&E), Prussian blue, and terminal deoxynucleotidyl transferase dUTP nick end labeling (TUNEL) with methyl green counterstain. Double-blind histological analyses were performed by a clinically qualified pathologist with expertise in sickle mice histology (see Supplementary Information for details).

### Statistics

Data are expressed as mean ± standard error of the mean (s.e.m.) unless otherwise specified, and the analyses were conducted using Prism software (GraphPad Prism, GraphPad Software, CA, USA) (see Supplementary Information for details).

## Results

### Short-term modulation of pain-related behaviors in wild-type and SCD mice

We hypothesized that tFUS stimulation applied to specific pain processing brain circuits could induce a significant change in thermal and mechanical sensitivity. To test this hypothesis, we administered single-session tFUS stimulation with a low PRF to S1HL in wild-type mice. Previous studies demonstrated that inhibitory neuronal activities can be evoked with a low PRF condition^36^. Furthermore, given the capability of the H276 device to specifically target one hemisphere of the mouse brain region, we evaluated the unilateral effect of tFUS on behavioral nocifensive responses to a noxious heat stimulus by comparing ipsilateral and contralateral withdrawal responses^35^. The delivery of ultrasound pulses with a PRF of 40 Hz to the S1HL in the left hemisphere of the brain resulted in a significant suppression of heat-pain related behaviors in both male (Fig. 3A) and female (Fig. 3B) wild-type mice. This suppression is represented as a marked change in the averaged Δ hPWL for 8-minute after tFUS compared to the baseline value, as well as sham treatment. However, Δ hPWL in the sham treated mice did not show a significant change compared to the pre-treatment baseline (Fig. 3A and B).

**Figure 3.**
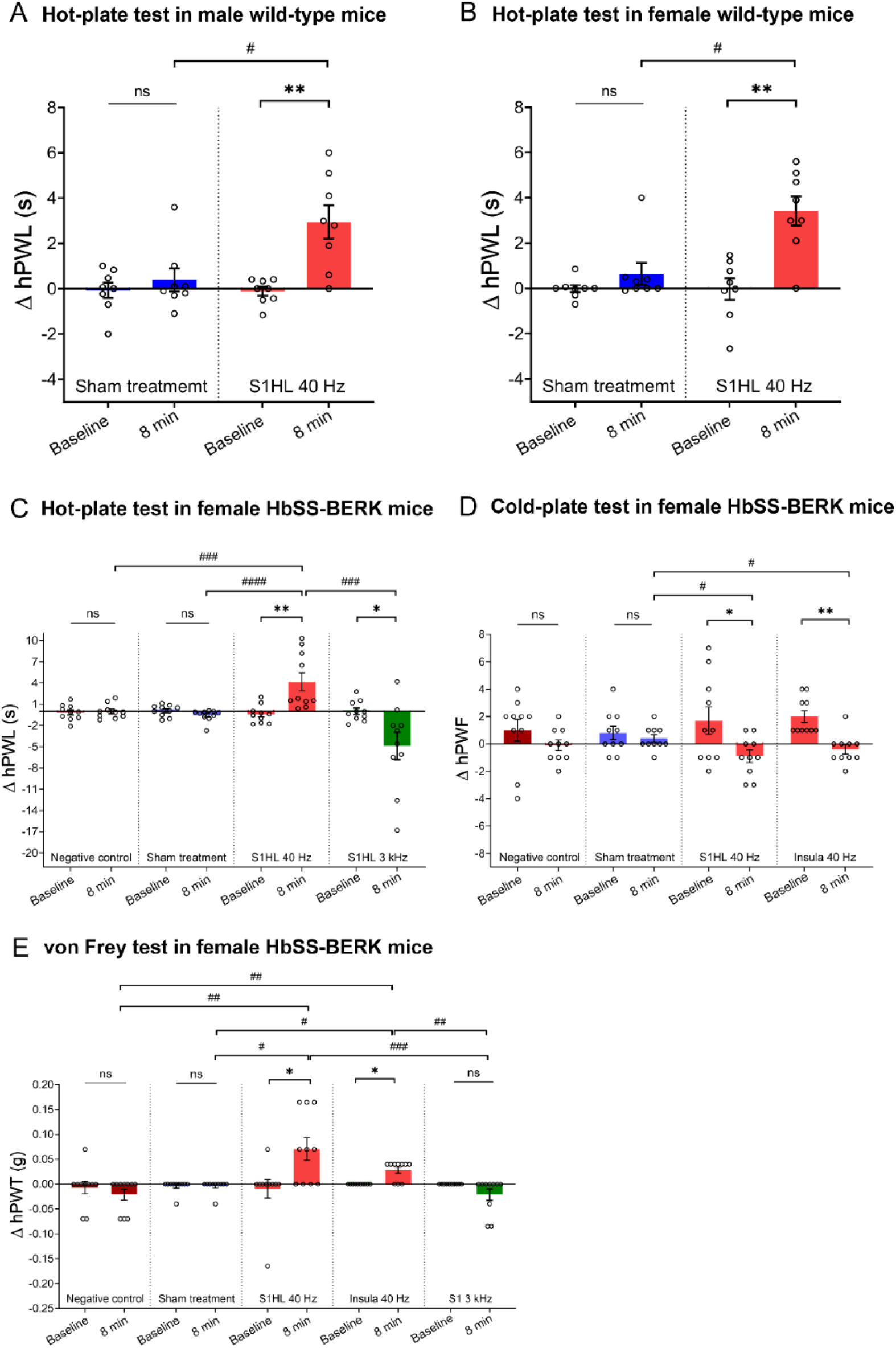
Short-term modulation of pain-related behaviors in wild-type mice and female HbSS-BERK mice. (A-B) Short-term inhibition of heat pain sensitivity in wild-type mice. The impact of tFUS on heat pain-associated behaviors was assessed in both (A) male (N=8) and (B) female (N=8) wild-type mice. Single-session tFUS with a PRF of 40 Hz at left S1HL resulted in a significant increase in latency difference between contralateral (right) and ipsilateral (left) hind paw compared to baseline and sham-treated mice. (C-E) Short-term modulation of thermal and mechanical hyperalgesia in female HbSS-BERK mice. Quantification of averaged Δ hPWL from hot-plate test, Δ hPWF from cold-plate test, and Δ hPWT from von Frey test was conducted before and after tFUS with controls groups (negative control and sham treatment) in HbSS-BERK mice. (C) Single-session tFUS with a PRF of 40 Hz at left S1HL (N=10) led to a significant increase in the averaged Δ hPWL from hot-plate test compared to baseline, sham treatment (N=10), and negative control (N=10). Control groups did not show apparent treatment effect compared to pre-treatment baseline. A negative value of Δ hPWL was observed with a PRF of 3 kHz at left S1HL (N=10), which was significantly different from the baseline and response with the PRF of 40 Hz. (D) Single-session tFUS with PRF of 40 Hz at left S1HL (N=10) and insula (N=10) resulted in a significant change in the Δ hPWF from cold-plate test at 8 min after treatment compared to the sham treatment and baseline. (E) tFUS with PRF of 40 Hz at left S1HL (N=10) and insula (N=10) led to a prominent elevation of Δ hPWT from von Frey test compared to sham-treated (N=10) and negative control mice (N=10). A lower withdrawl threshold of the contralateral (right) hind paw was observed with tFUS at PRF of 3 kHz at left S1HL(N=10) compared to PRF of 40 Hz at S1HL and insula. ns: not significant, \**p*<0.05, \*\**p*<0.01 using t-test with Wilcoxon matched-pairs signed rank test; ^#^*p*<0.05, ^##^*p*<0.01, ^###^*p*<0.001, ^####^*p*<0.0001 using t-test with Mann-Whitney test.

We subsequently investigated the impact of tFUS stimulation on the well-established humanized mouse model of SCD (HbSS-BERK) that constitutively displays significant thermal and mechanical hyperalgesia, which is similar to that observed in human SCD patients^28,37^. Given that higher pain intensity has been reported in female patients and female HbSS-BERK mice compared to males and Townes sickle mice^28,32^, testing a novel pain treatment for females is of importance and priority. We evaluated responses to heat stimuli by comparing ipsilateral (left) and contralateral (right) hPWL values modulated by tFUS stimulation or control groups in female HbSS-BERK mice (Fig. 3C and Supplementary Fig. S4A). Our findings indicated that a single-session tFUS with a PRF of 40 Hz applied to the left S1HL resulted in a remarkable suppression of heat hypersensitivity-related behaviors. This suppression was evident through a significant change in the averaged Δ hPWL for 8-minute after tFUS, compared to the baseline value as well as negative control and sham treatment. However, the Δ hPWL in the control groups is similar to the pre-treatment baseline. We also observed a negative value of Δ hPWL stimulated by the single-session tFUS with a PRF of 3 kHz to the left S1HL, indicating an increased contralateral hind paw hypersensitivity. In light of these distinct behavioral responses to tFUS stimulation paradigms, a bidirectional modulation of heat hypersensitivity-related behaviors was further examined by adjusting PRF values to generate attenuation or exacerbation of heat hyperalgesia in female HbSS-BERK mice (Supplementary Fig. S8B).

Subsequently, we investigated whether single-session tFUS stimulation could modulate cold hyperalgesia in female HbSS-BERK mice by comparing ipsilateral and contralateral hPWF values during a testing period of 2-minute (Fig. 3D and Supplementary Fig. S4B). Compared to the pre-application baseline, single-session tFUS stimulation with a 40 Hz PRF to the left S1HL or insula led to a significant decrease in cold hyperalgesia, represented as a negative value of Δ hPWF. The effect of single-session tFUS stimulation at the left S1HL and insula on Δ hPWF significantly differed from the sham group at 8-minute after treatment but not from the control groups, tFUS with a PRF of 40 Hz at the thalamus, and tFUS with a PRF of 3 kHz (Supplementary Fig. S3B).

Additionally, we investigated how single-session tFUS stimulation modulates mechanical hyperalgesia in HbSS-BERK mice by comparing the withdrawal threshold of the ipsilateral (left) hind paw with that of the contralateral (right) hind paw (Fig. 3E and Supplementary Fig. S4C). When the left S1HL or insula, but not the thalamus, was stimulated with tFUS at a PRF of 40 Hz, a significant reduction in mechanical hyperalgesia, indicated by Δ hPWT, was observed compared to the baseline as well as sham-treated or negative control mice. Similar to the observed behavior changes in the heat hyperalgesia (Fig. 3C), tFUS with a PRF of 3 kHz at the left S1HL also exacerbated mechanical hypersensitivity, represented by a lower withdrawal threshold of the contralateral (right) hind paw compared to the ipsilateral (left) one.

### Sustained amelioration of heat hyperalgesia in sickle mice with tFUS

We investigated achieving a sustained tFUS effect on heat hyperalgesia by targeting deeper brain structures including insula and thalamus. The single-session tFUS with the 40 Hz PRF at the left insula showed a much longer latency of the contralateral hind paw compared to the ipsilateral hind paw for 30 minutes, which differed from the pre-stimulation baseline. This effect was specific to the tFUS stimulation, as neither control groups (negative control and sham treatment) nor tFUS with the 3 kHz PRF significantly differed from their respective baseline values (Fig. 4A). Significant differences were also observed between tFUS with the 40 Hz PRF and controls. In contrast, delivering tFUS with the 40 Hz PRF to the thalamus resulted in a less pronounced behavioral response with a tendency towards attenuating heat pain-related behaviors (Fig. 4B). Collectively, single-session tFUS stimulation with the 40 Hz PRF to the insula, but not the thalamus, is effective for a sustained attenuation (30-minute ≤ effect <1-hour) of hyperalgesia to heat stimuli in female HbSS-BERK mice.

We further examined whether a sustained attenuation (≥1-hour) of pain behaviors could be achieved in older female sickle mice, as treatment for chronic pain in older adults is a major challenge^28,37,38^. We observed a significant change in the behavioral response to heat stimulus, represented as the averaged hPWL, in older female HbSS-BERK mice (10-15 month-old) compared to those in younger female mice (5-9 month-old) (Supplementary Fig. S5A). Given significantly higher heat hyperalgesia in older female HbSS-BERK mice, we administered single-session tFUS to S1HL for 20-minute and for 1-hour, and as a result, only 1-hour of stimulation was found to produce a short-term suppression of heat hyperalgesia in older female HbSS-BERK mice (Supplementary Fig. S5B). Thus, to achieve a sustained effect in older female HbSS-BERK mice, we administered multi-session tFUS stimulation to the selected brain regions for 14 days, with 1-hour dose per day, on the behavioral responses to noxious heat stimuli (Fig. 4C). On Day 14, 2 hours after the 1-hour treatment, we observed a significantly longer latency of the contralateral hind paw withdrawal than that of the ipsilateral hind paw response when compared with pre-stimulation baseline or sham-treated group (Fig. 4D). In addition, the effect of multi-session tFUS, but not sham treatment, lasted at least 2 hours, but less than 4 hours according to extensive behavioral assessments at 2-, 4-, and 6-hour after the 14-day treatment (Supplementary Fig. S5C and D). Together, these results suggest that 1-hour tFUS daily stimulation for 14 days with a PRF of 40 Hz at S1HL or insula can induce a sustained amelioration (2-hour ≤ effect <4-hour) of heat hyperalgesia in older female HbSS-BERK mice.

**Figure 4.**
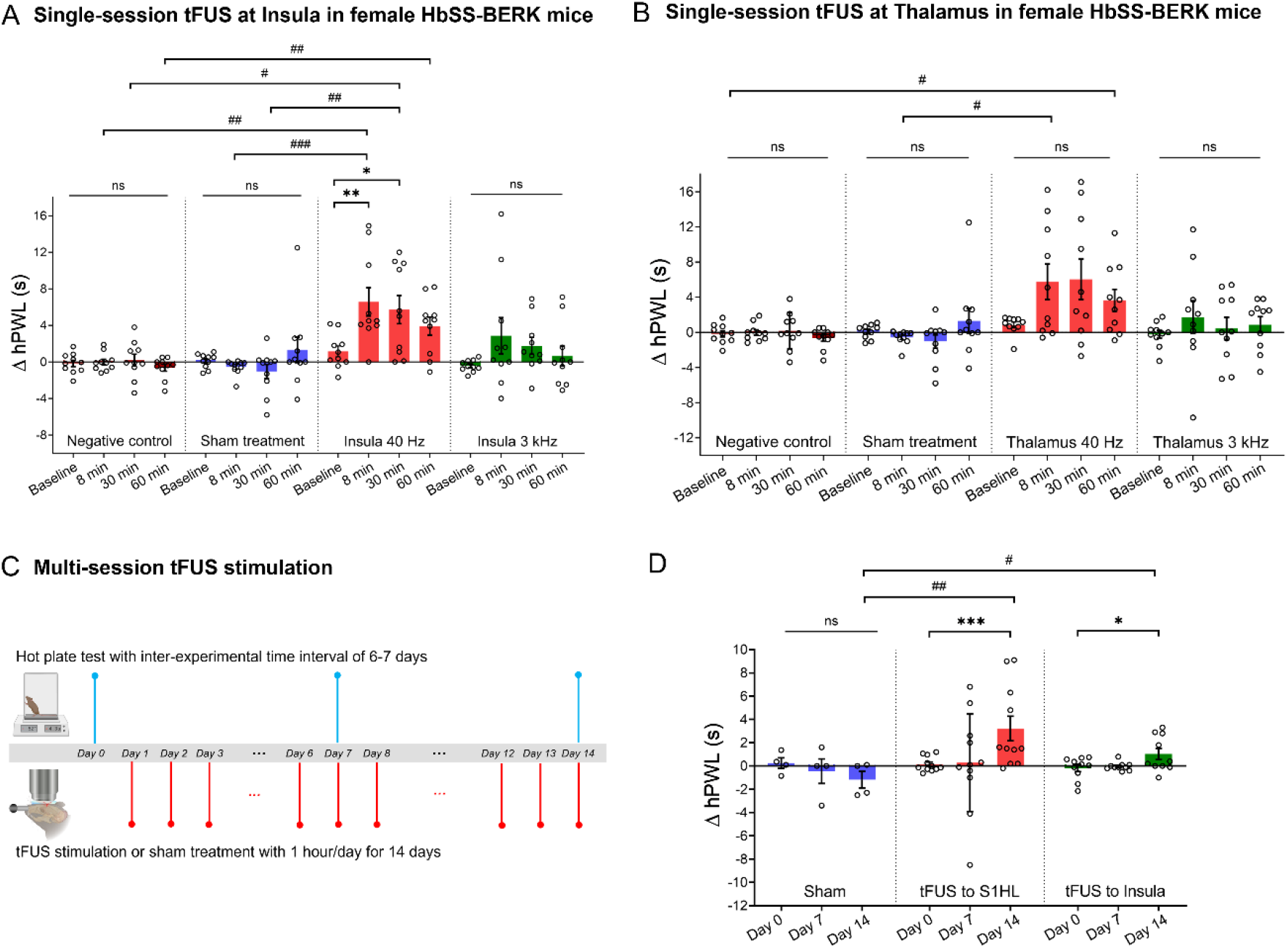
Sustained modulatory effect on heat hyperalgesia with the single- and multi-session tFUS stimulation in female HbSS-BERK mice. (A) The averaged Δ hPWL stimulated by single-session tFUS with a PRF of 40 Hz at left insula (N=10) was significantly increased up to 30 min in HbSS-BERK mice compared to the pre-tFUS baseline, as well as values from sham (N=10) and negative control groups (N=10). Single-session tFUS with a PRF of 3 kHz (N=10), sham treatment, and negative control did not result in remarkable changes in Δ hPWL compared to the baseline. (B) Targeting the thalamus with tFUS using a PRF of 40 Hz (N=10) and 3 kHz (N=10) did not result in a prominent modulatory effect compared to the pre-stimulation baseline, as well as sham-treated mice (N=10) and negative control mice (N=10), except for two measurements (right panel, ^#^*p*=0.0104, 8 min after tFUS with the PRF of 40 Hz and sham treatment; ^#^*p*=0.0277, 60 min after tFUS with the PRF of 40 Hz and negative control). (C) Sustained amelioration of heat hyperalgesia was examined in older female HbSS-BERK mice (10 to 15 month-old) by measuring latency to noxious heat stimuli compared to pre-stimulation baseline upon 14-day tFUS stimulation. (D) After multi-session tFUS stimulation with a PRF of 40 Hz to S1HL (N=11) or insula (N=10) for 14 days, the averaged changes of Δ hPWL were significantly increased for at least 2 h compared to the baseline values, as well as sham-treated behavioral change (N=4). The measurement was conducted at 2 h after tFUS or sham treatment, indicating that the effect of multi-session tFUS stimulation is considered to persist for at least 2 h. ns: not significant, \**p*<0.05, \*\**p*<0.01, \*\*\**p*<0.001 using one-way ANOVA with Friedman test or t-test with Wilcoxon one-tailed signed rank test; ^#^*p*<0.05, ^##^*p*<0.01, ^###^*p*<0.001 using one-way ANOVA with Kruskal-Wallis test or t-test with one-tailed Mann-Whitney test.

### Electrophysiological evaluation of heat stimulus-evoked brain activity and modulation of pain with tFUS

To better understand how tFUS stimulation modulates neuronal activities in the targeted brain circuits, we quantified chronic pain based on a significant broadband increase in neuronal oscillations, spanning theta, alpha, and beta band activities in left and right S1HL (Supplementary Fig. S6A) of female HbSS-BERK mice compared to the power in female wild-type mice (Fig. 5A). This increase represents a key feature of chronic pain condition in the humanized HbSS-BERK mice and is consistent with prior animal studies^39,40^. Furthermore, we found that high-temperature hind paw stimulation (HTHPS, a noxious stimulus), but not low-temperature hind paw stimulation (LTHPS, an innocuous stimulus), induced a significant decrease in theta, alpha, and beta power in the contralateral S1HL (Supplementary Fig. S6B) in HbSS-BERK mice compared to baseline responses (Fig. 5B). This finding is consistent with previous experimental observations^41,42^. Leveraging the EEG pain indicators, we studied the effect of tFUS stimulation at the right S1HL on the brain rhythms underlying painful heat stimuli delivered to the plantar surface of the contralateral hind paw. After single-session tFUS stimulation with a PRF of 40 Hz, a significant increase in Δ frequency power between HTHPS and LTHPS (i.e., normalized power in HTHPS - averaged normalized power in LTHPS) was observed (Fig. 5C and Supplementary Fig. S6C-F). This distinct brain electrophysiological response to tFUS stimulation with a PRF of 40 Hz supports the observed amelioration of heat pain-related behaviors.

**Figure 5.**
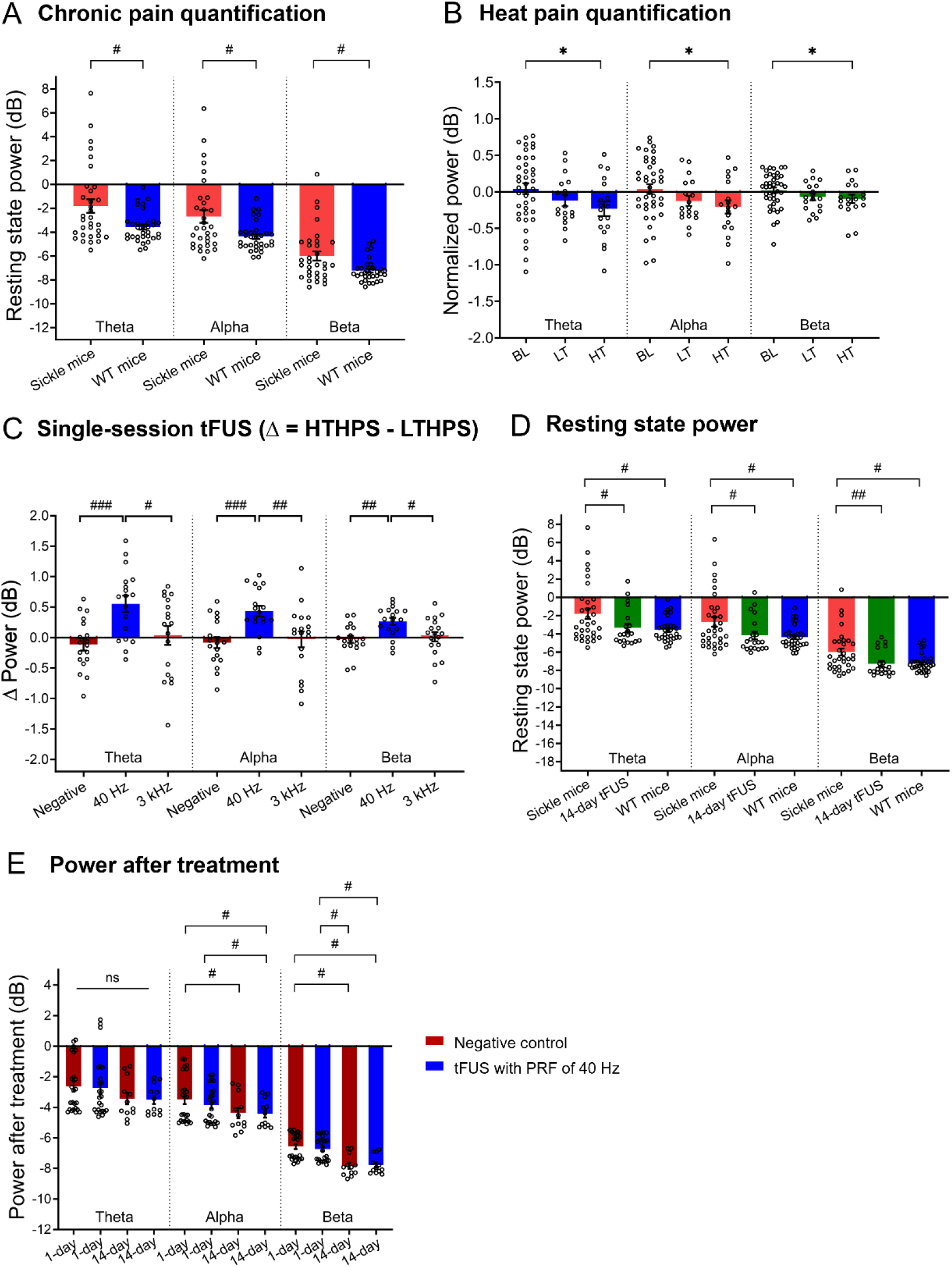
Modulation of heat stimulus-evoked brain activity and chronic pain resting state in female HbSS-BERK mice. (A) Chronic pain quantification in brain rhythms at rest, showing a broadband increase of oscillations from theta to beta frequencies in female HbSS-BERK mice (N=8) compared to female wild-type mice (N=8). (B) Heat pain quantification in brain oscillations from the negative control group. High-temperature hind paw stimulation (HTHPS), but not low-temperature hind paw stimulation (LTHPS), resulted in a significant decrease in theta, alpha, and beta power in the contralateral brain region of S1HL in HbSS-BERK mice (N=8). Note that baseline (BL), low temperature (LT) of 26.2 ± 0.2°c < T ≤ 36.2 ± 0.2°c, and high temperature (HT) of 38.9 ± 0.2°c < T ≤ 46.4 ± 0.1°c. (C) Differences (Δ) of oscillatory power during normalized HTHPS and averaged normalized LTHPS. Post-tFUS with a PRF of 40 Hz, but not 3 kHz, at S1HL induced a significant increase in Δ power from theta to beta frequencies (N=8), contrasting the heat pain-related EEG pattern. (D) Multi-session tFUS at S1HL induced a significant decrease in power from theta to beta frequencies (N=5), showing a remarkable difference from the power of untreated HbSS-BERK mice while being similar to the power of wild-type mice. (E) At 80 s following heat stimulus at hind paw, power in alpha and beta frequencies was significantly suppressed by multi-session tFUS with a PRF of 40 Hz but not by single-session tFUS with a PRF of 40 Hz (N=8 for single-session tFUS and N=4 for multi-session tFUS). Ns: not significant, \**p*<0.05 using t-test with Wilcoxon signed rank test; ^#^*p*<0.05, ^##^*p*<0.01, ^###^*p*<0.001 using t-test with Mann-Whitney test.

Furthermore, we extensively examined whether multi-session tFUS affects the resting-state EEG power in older female HbSS-BERK mice, which demonstrated sustained suppression of pain behaviors. We measured the resting-state EEG power on the day following 14-day repeated tFUS stimulation (1-hour tFUS treatment per day) and found a significant broadband decrease in neuronal oscillatory activities spanning from theta to beta frequencies in the left and right S1HL region compared to the power in female HbSS-BERK mice (Fig. 5D). This decrease brought the oscillatory powers close to those observed in wild-type mice. Additionally, we examined the impact of post-tFUS stimulation following heat stimuli and found a significant power suppression in alpha and beta frequencies compared with multi- and single-session tFUS with a PRF of 40 Hz in post-time periods of 6-minute (Fig. 5E).

### tFUS safety evaluation

We tested whether tFUS stimulation to superficial and deep brain regions would negatively impact normal motor coordination^43,44^. After the single-session tFUS at S1HL or thalamus and multi-session tFUS stimulation to S1HL or insula, we did not find any significant difference in the averaged latency to fall on the rotarod compared with pre-stimulation baseline or negative control or sham treatment (Fig. 6A and B). Similarly, the average time spent on the rotarod from the negative control, or the multi-session sham treatment was not markedly different from the baseline values, indicating that single- and multi-session tFUS stimulation at cortical and deep brain structures does not induce a remarkable adverse effect on motor function in HbSS-BERK mice.

**Figure 6.**
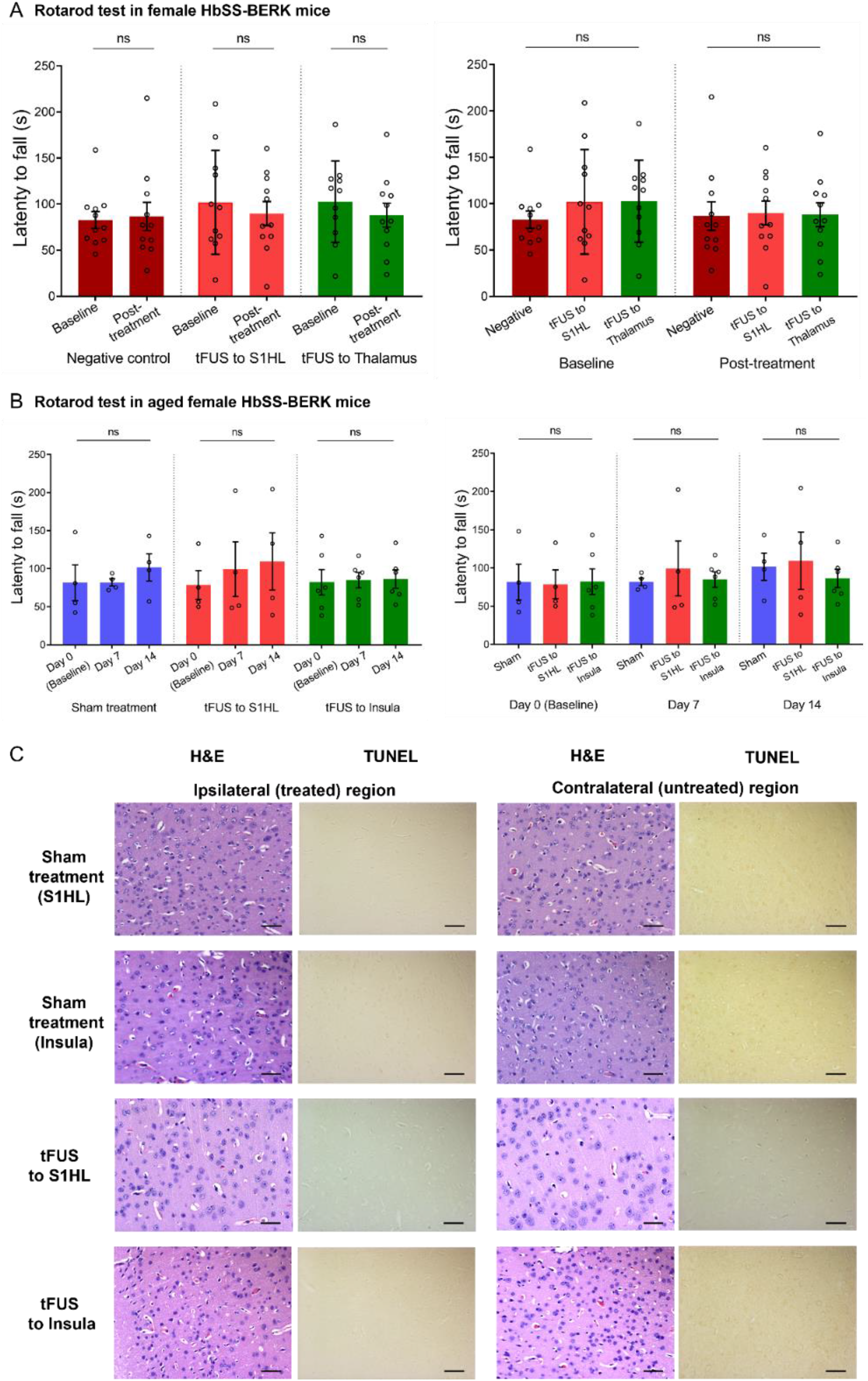
tFUS safety evaluation in female HbSS-BERK mice. (A-B) The effect of tFUS on the motor coordination and balance was investigated in rotarod testing. After 60 min of single-session tFUS stimulation with a PRF of 40 Hz to S1HL (N=11) or thalamus (N=11) and after 60 min of multi-session tFUS with a PRF of 40 Hz to S1HL (N=4) or insula (N=6) for 14 days with 1 h per day, no significant alteration was observed in the averaged latency to fall compared with pre-stimulation baseline as well as control groups. Additionally, there was no significant difference of control groups from their baseline values. ns: not significant using t-test with Wilcoxon matched-pairs signed rank test and one-way ANOVA with Kruskal-Wallis test. (C) Double-blind histological analysis using H&E and TUNEL stains showed that multi-session tFUS stimulation to the left S1HL or insula had no adverse impact on tissue structural integrity and nuclear morphology within the treated brain region compared with contralateral brain regions and sham treatment. Scale bars: 50 µm.

To further examine the possibility of adverse effects of tFUS stimulation, a double-blind evaluation of brain histology was performed after chronic treatment (1-hour per day over 14 days) in tFUS and control groups of HbSS-BERK mice (Fig. 6C) and genetic control mice, HbAA-BERK (Supplementary Fig. S7). We did not observe any significant change in the structural integrity of the tissues, iron deposits, and the number of apoptotic cells in the ipsilateral (treated) and contralateral (untreated) brains as well as those from the sham, negative control, and multi-session high-dose tFUS treatment (Fig. 6C and Supplementary Fig. S7).

## Discussion

We have developed a novel non-invasive and non-pharmacologic tFUS neuromodulation approach as an analgesic intervention without causing any adverse effects on neuropathology or brain function in mice. We demonstrated that single- and multi-session tFUS stimulation non-invasively and distinctly interacts with pain processing circuits in the brain, leading to effective modulation of pain-associated behaviors in wild-type and SCD mice. We additionally observed significant changes in electrophysiological activities related to pain along with tFUS neuromodulation.

Previous studies have demonstrated that optogenetic activation of parvalbumin (PV) neurons at 40 Hz or tFUS at 40 Hz in the hippocampus or S1 triggers activation of immune cells^45–47^ or positively impacts mechanical nociception^48^, respectively. In contrast, vibroacoustic treatment at 40 Hz leads to a beneficial effect on chronic pain disorders such as fibromyalgia with widespread musculoskeletal pain^49,50^. We observed that repeated ultrasound pulses at 40 Hz to multiple brain regions exhibit a distinct contrast to the brain oscillatory pattern of pain and ameliorate pain-associated behaviors. Our findings suggest that the inhibitory effects of tFUS using a PRF of 40 Hz at the S1HL can modulate cortical processes of pain-related sensory signals from peripheral input, reducing its function in pinpointing the exact location of pain on the body and differentiating the type of pain stimuli, akin to numbing the peripheral tissue. In addition to the effect of specifically repeated pulse rates, a lower duty cycle has been shown to induce inhibitory effects in cortical and thalamic neuronal activity^51,52^. Thus, we further investigated whether tFUS with specific PRF values within a low duty cycle, while maintaining the same transmitted ultrasound energy, could lead to a reduction in sensitivity to a nociceptive stimulus. Neither tFUS stimulation with additional parameter 1 (AP1: PRF of 10 Hz and TBD of 800 µs) nor additional parameter 2 (AP2: PRF of 3 kHz and TBD of 2.67 µs) resulted in a significant change in heat pain-related behaviors in HbSS-BERK mice compared to pre-stimulation (Supplementary Fig. S8A). In addition, we adjusted PRF values from 40 Hz to 3 kHz and found that tFUS with a PRF of 1 kHz can be considered as a turning point from generating neural inhibition (suppressing pain) to inducing neural excitation (intensifying pain) when tFUS is applied to S1HL (Supplementary Fig. S8B). Therefore, the ultrasound PRF value plays a key role in modulating behavioral responses to noxious stimuli in HbSS-BERK mice.

We also found a remarkable change in brain oscillations stimulated by the single-session tFUS with PRF of 3 kHz to the S1HL from the innocuous LTHPS period (Supplementary Fig. S6C-F), which may be induced by facilitated cortical excitability^53^ built up during the excitatory tFUS neuromodulation (PRF: 3 kHz) (Supplementary Fig. S6G-I). This finding is consistent with prior studies demonstrating critical contribution of CaMKIIα on chronic pain exaggeration in HbSS-BERK mice^30,54^, increased power in low frequency bands by optogenetically suppressing e.g., cortical PV inhibitory neurons^53^, and tFUS-mediated distinct action potential waveforms and significant spike rates of S1 in transgenic CaMKIIα mice and PV mice^36^.

We conducted a comprehensive set of control experiments to evaluate whether the achieved analgesic effect is specific to tFUS stimulation. Firstly, to prevent the potential analgesic effect of anesthesia, we used an awake experimental setup (i.e., a head-restrained awake naturally walking mouse in the air-lifted platform) to directly measure the tFUS modulatory effect (Supplementary Fig. S9A-B). As we also observed a consistent significant tFUS modulatory effect in head-fixed awake mice, the light anesthesia used in this study does not confound the conclusion that tFUS can modulate pain-associated behavioral responses. Secondly, considering the possibility of altered nocifensive behaviors in the repeated hot-plate test due to non-nociceptive factors such as learning or habituation^55,56^, we examined the effect of inter-experimental day intervals to rule out the potential contribution of non-nociceptive factors to the observed tFUS-mediated analgesic effect (Supplementary Fig. S9C). After measuring latency from noxious heat stimuli at every 3-4 days (but not 6-7 days), we observed a significant decrease in the hPWL from the measurement on Day 10-11 compared to the hPWL on Day 1 in HbSS-BERK mice. This suggests that an inter-experimental time interval of 6-7 days can minimize the impact of non-nociceptive factors on the pain-related behaviors modulated by tFUS.

For clinical translation, low-intensity ultrasound (I_SPTA_: 208.46 mW/cm^2^ at the highest PRF of 3 kHz and I_SPPA_: 0.35 W/cm^2^ at the TBD of 200 μs measured behind a freshly excised female mouse skull sample) was used to adhere to the regulatory standards for diagnostic ultrasound safety (I_SPTA_ ≤ 720 mW/cm^2^, I_SPPA_ ≤ 190 W/cm^2^) set by the Food and Drug Administration (FDA)^57,58^. We further investigated the impact of tFUS on heat pain-associated behaviors in female HbAA-BERK control and HbSS-Townes mice. Notably, single-session tFUS with a PRF of 40 Hz significantly suppressed heat pain-related behaviors in female HbAA-BERK control mice (Supplementary Fig. S10A) and female HbSS-Townes mice (Supplementary Fig. 10B). In particular, with different globin chains^32^ and fetal hemoglobin levels closely related to patterns of pronociceptive and antinociceptive brain connectivity in Townes sickle mice^59^, this may contribute to more gradual tFUS effects on pain behaviors compared to the more spontaneous tFUS effects observed in BERK mice. However, the efficacy of single-session tFUS with a PRF of 40 Hz to S1HL or insula was notably diminished in older female and male HbSS-BERK mice (Supplementary Fig. S11A-D), as well as older female wild-type mice treated by tFUS to S1HL, given their remarkably increased heat pain sensitivity (Supplementary Fig. S11E and F). These observations highlight the need to establish a patient-specific dosage window for tFUS stimulation, primarily based on the age of chronic pain patients. Notably, this dosage window seems to be less dependent on gender differences (Supplementary Fig. S11A-D) compared to age. Based on the histological investigations, we established an extended safety window for tFUS stimulation using high-dose tFUS stimulation (transcranial peak-peak pressure: 2059.75 kPa, I_SPTA_: 262.30 mW/cm^2^ at the PRF of 40 Hz and I_SPPA_: 32.79 W/cm^2^ at the TBD of 200 μs measured behind a freshly excised female mouse skull sample) to prevent short- and long-term adverse effects.

From a neuromodulation perspective, the promising evidence of sustained suppression of heat pain-associated behaviors, coupled with a significant change in the resting-state power after repeated stimulations for 14-day, underscore the need for a deeper understanding of how multi-session tFUS interacts with cellular and physiological functions of pain processing brain circuits. The presented results not only merit further investigation into the efficacy of multi-session tFUS beyond the 14-day period but also call for neurobiological assessments of the induced long-lasting analgesic effects. Moreover, exploring the impact of simultaneously targeting multiple pain processing brain circuits, e.g., co-stimulations at S1 and insula, may reveal the potential for generating more amplified inhibitory effects on pain-associated behaviors than respective S1 or insula stimulation alone.

The preclinical results presented here showcase a unique bidirectional modulation of pain-associated behaviors through tailored tFUS stimulation, employing both effective and ineffective PRF. This connects fundamental neuroscience knowledge of tFUS neuromodulation to behaviors and brain oscillations observed in HbSS-BERK mice. Specifically, inhibitory tFUS stimulation (at 40 Hz PRF) significantly alleviates pain by inhibiting neuronal activities at a specific brain target. Conversely, excitatory tFUS stimulation (at 3 kHz PRF) amplifies the pain signaling through neuronal excitation at the same brain location. The distinct outcomes expected from applying different PRF levels can be explained by considering specific neuronal oscillations in the pain processing brain circuit, underlying responses to innocuous and painful stimuli, and our previous study on the intrinsic functional neuron-type selectivity of tFUS neuromodulation^36^. To translate this technology for treating chronic pain in humans, applying the ultrasound PRF of 40 Hz for inhibitory effects is recommended. However, it is crucial to note that modifying the ultrasound fundamental frequency and increasing ultrasound pressure may be required to efficiently deliver ultrasound energy through thick human skulls for modulating larger human brain targets effectively.

In summary, our study marks the first demonstration of the significant impact of non-invasive low-intensity transcranial focused ultrasound (tFUS) stimulation on cortical and deep brain targets associated with pain processing. This stimulation resulted in a significant change in brain oscillations and modulation of thermal and mechanical pain-associated behaviors in *in vivo* mouse models. Our findings contribute to a fundamental understanding of bidirectional modulation in neuronal oscillations and pain-associated behaviors, emphasizing the effects of pulse repetition frequency (PRF) and highlighting sustained pain suppression achievable through deeper cortical structure targeting and multi-session tFUS stimulation. The experimental evidence presented, coupled with the significant advancements in the efficacy and safety of tFUS neuromodulation using a state-of-the-art multi-element highly focused random array transducer, underscores the translational potential of tFUS as an effective, non-pharmacological, and non-invasive device-based neuromodulation technique for managing chronic pain.

## Acknowledgments

This work was supported in part by NIH EB029354, NS124564, NS131069, and HL147562. S.K. was supported by Diversity Supplement 3R01HL147562-03S, and D.A. was supported by a University of California President’s Fellowship and Giannini Foundation Fellowship. *The content is solely the responsibility of the authors and does not necessarily represent the official views of the National Institutes of Health*.

The authors would like to thank Natalie Garcia and Reina Lomelli, University of California Irvine for their availability of transgenic mice. We also thank Kyle Morrison from Sonic Concepts, Inc. for providing consultation in customizing the H276 probe. Some cartoons in Figures and Supplementary Figures were created with BioRender.com.

## Author Contributions

B.H., K.Y. initiated the concept; B.H., K.Y., M.G.K., K.G. designed the studies; M.G.K. conducted behavior experiments with neuromodulation and analyzed behavior data; M.G.K., C-Y.Y. performed EEG experiments with neuromodulation and analyzed electrophysiological data; K.Y., Y.N., C-Y.Y., D.A. conducted behavior data assessments; K.G., D.A., S.A. produced sickle cell mice; K.Y., K.K., Q.C. designed the multi-element array transducer; K.Y. simulated the design of random array ultrasound transducer in experimental setup; K.Y., M.G.K., X.N. characterized the transducer array and validated the ultrasound targeting; R.F., D.A. performed histology assessments; M.G.K., K.Y. co-wrote the original draft; B.H., K.G., R.F., D.A., C-Y.Y., K.K., Q.Y., Y.N. contributed to reviewing and editing of the manuscript. B.H. and K.G. supervised the work.

## Data Availability

Data supporting the conclusions are presented in the paper and supplementary information. Additional data on EEG recordings and behavior results will be shared in a public repository upon paper acceptance.

---

**Table 1.**
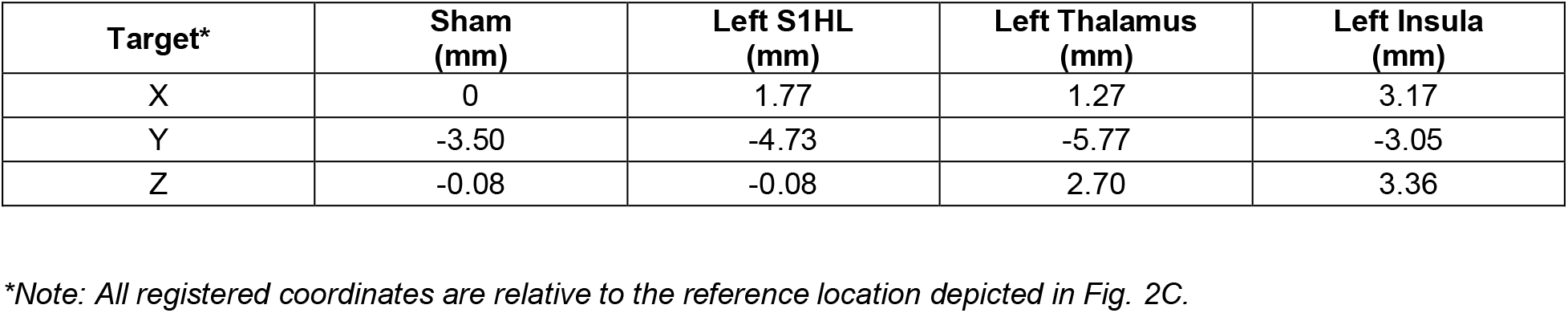
Registered coordinates of ultrasound focus on respective brain targets based on Allen Mouse Brain Atlas.

## References

1. Goldberg DS, McGee SJ. Pain as a global public health priority. BMC Public Health 2011;11:770.

2. Pitcher MH, Von Korff M, Bushnell MC, Porter L. Prevalence and profile of high impact chronic pain in the United States. Journal of Pain 2019;20(2):146–160.

3. Okie S. A flood of opioids, a rising tide of deaths. N. Engl. J. Med. 2010;363(21):1981–1985.

4. Ballantyne JC. Opioids for the treatment of chronic pain: mistakes made, lessons learned, and future directions. Anesth. Analg. 2017;125(5):1769–1778.

5. Boccard SG, Pereira EA, Aziz TZ. Deep brain stimulation for chronic pain. J. Clin Neurosci. 2015;22(10):1537–1543.

6. Lefaucheur JP, Drouot X, Keravel Y, Nguyen JP. Pain relief induced by repetitive transcranial magnetic stimulation of precentral cortex. Neuroreport. 2001;12(13):2963–5.

7. Knotkova H, Greenberg A, Leuschner Z, Soto E, Cruciani RA. Evaluation of outcomes from transcranial direct current stimulation (tDCS) for the treatment of chronic pain. Clinical Neurophysiology. 2013;12(10):e125–6.

8. Johnson MD, Lim HH, Netoff TI, et al. Neuromodulation for brain disorders: challenges and opportunities. IEEE Trans. Biomed Eng. 2013;60(3):610–624.

9. Wagner T, Valero-Cabre A, Pascual-Leone A. Noninvasive human brain stimulation. Annu. Rev. Biomed. Eng. 2007;9:527–565.

10. Fry FJ, Ades HW, Fry WJ. Production of reversible changes in the central nervous system by ultrasound. Science. 1958;127(3289):83-4.

11. Tufail Y, Yoshihiro A, Pati S, Tauchmann ML, Tyler WJ. Ultrasonic neuromodulation by brain stimulation with transcranial ultrasound. Nat. Protoc. 2011;6(9):1453–1470.

12. Yoo SS, Bystritsky A, Lee JH, et al. Focused ultrasound modulates region-specific brain activity. Neuroimage. 2011;56(3):1267–1275.

13. Kubanek J. Neuromodulation with transcranial focused ultrasound. Neurosurg Focus 2018;44(2):E14.

14. Verhagen L, Gallea C, Folloni D, et al. Offline impact of transcranial focused ultrasound on cortical activation in primates. eLife 2019;8:e40541.

15. Yang Y, Yuan J, Field RL, et al. Induction of a torpor-like hypothermic and hypometabolic state in rodents by ultrasound. Nat Metab 2023;5(5):789-803.

16. Wang Z, Yan J, Wang X, Yuan Y, Li X. Transcranial ultrasound stimulation directly influences the cortical excitability of the motor cortex in parkinsonian mice. Mov Disord. 2020;35(4):693–698.

17. Fasano A, Vloo PD, Llinas M, et al. Magnetic resonance imaging-guided focused ultrasound thalamotomy in Parkinson tremor: reoperation after benefit decay. Mov Disord. 2018;33(5):848–9.

18. Beisteiner R, Matt E, Fan C, et al. Transcranial pulse stimulation with ultrasound in Alzheimer’s disease-a new navigated focal brain therapy. Adv Sci. 2019;7(3):1902583.

19. Badran BW, Caulfield KA, Stomberg-Firestein S, et al. Sonication of the anterior thalamus with MRI-guided transcranial focused ultrasound, (tFUS) alters pain thresholds in healthy adults: a double-blind, sham-controlled study. Brain Stimul. 2020;13(6):1805–12.

20. Hameroff S, Trakas M, Duffield C, et al. Transcranial ultrasound (TUS) effects on mental states: a pilot study. Brain Stimul. 2013;6(3),409–15.

21. Chen LM, Yang PF, Newton A, et al. MRI guided focused ultrasound modulation of deep brain pain regions and circuits in nonhuman primates. Brain Stimul. 2021;14(6):1599.

22. Zhang T, Wang Z, Liang H, et al. Transcranial focused ultrasound stimulation of periaqueductal gray for analgesia. IEEE Trans Biomed Eng. 2022;69(10):3155–62.

23. Bushnell MC, Ceko M, Low LA. Cognitive and emotional control of pain and its disruption in chronic pain. Nat Rev Neurosci. 2013;14(7):502–11.

24. Peirs C, Seal RP. Neural circuits for pain: recent advances and current views. Science. 2016;354(6313):578-584.

25. Mogil JS. Animal models of pain: progress and challenges. Nat. Rev. Neurosci. 2009;10(4):283–94.

26. Deuis JR, Dvorakova LS, Vetter I. Methods used to evaluate pain behaviors in rodents. Front Mol Neurosci. 2017;10:284.

27. Pászty C, Brion CM, Manci E, Witkowska HE, Stevens ME, Mohandas N, Rubin EM. Transgenic knockout mice with exclusively human sickle hemoglobin and sickle cell disease. Science. 1997;278(5339):876-8.

28. Kohli DR, Li Y, Khasabov SG, et al. Pain-related behaviors and neurochemical alterations in mice expressing sickle hemoglobin: modulation by cannabinoids. Blood. 2010;116(3):456–65.

29. Cain DM, Vang D, Simone DA, Hebbel RP, Gupta K. Mouse models for studying pain in sickle disease: effects of strain, age, and acuteness. Br J Haematol. 2012;156(4): 535–544.

30. Tran H, Gupta M, Gupta K. Targeting novel mechanisms of pain in sickle cell disease. Blood. 2017;130(22):2377–2385.

31. Sagi V, Song-Naba WL, Benson BA, Joshi SS, Gupta K. Mouse models of pain in sickle cell disease. Curr Protoc Neurosci. 2018;85(1):e54.

32. Lei J, Benson B, Tran H, Ofori-Acquah SF, Gupta K. Comparative analysis of pain behaviours in humanized mouse models of sickle cell anemia. PLoS One. 2016;11(8): e0160608.

33. Bonin RP, Bories C, De Koninck YA simplified up-down method (SUDO) for measuring mechanical nociception in rodents using von Frey filaments. Mol Pain. 2014;10:26.

34. Vincent L, Vang D, Nguyen J, Benson B, Lei J, Gupta K. Cannabinoid receptor-specific mechanisms to alleviate pain in sickle cell anemia via inhibition of mast cell activation and neurogenic inflammation. Haematology. 2015;101(5):566–577.

35. Lavich TR, Cordeiro RS, Silva PM, Martins MA. A novel hot-plate test sensitive to hyperalgesic stimuli and non-opioid analgesics. Braz J Med Biol Res. 2005;38(3):445–451.

36. Yu K, Niu X, Krook-Magnuson E, He B. Intrinsic functional neuron-type selectivity of transcranial focused ultrasound neuromodulation. Nat Commun. 2021;12:2519.

37. Hillery CA, Kerstein P, Vilceanu D, et al. Transient receptor potential vanilloid 1 mediates pain in mice with severe sickle cell disease. Blood. 2011;118(12):3376–83.

38. García-Esquinas E, Rodríguez-Sánchez I, Ortolá R, et al. Gender differences in pain risk in old age: magnitude and contributors. Mayo Clin Proc. 2019;94(9):1707–1717.

39. LeBlanc BW, Bowary PM, Chao YC, Lii TR, Saab CY. Electroencephalographic signatures of pain and analgesia in rats. Pain 2016;157(10):2330–2340.

40. LeBlanc BW, Lii TR, Huang JJ, et al. T-type calcium channel blocker Z944 restores cortical synchrony and thalamocortical connectivity in a rat model of neuropathic pain. Pain 2016;157(1):255–264.

41. Peng W, Hu L, Zhang Z, Hu Y. Changes of spontaneous oscillatory activity to tonic heat pain. PLoS One 2014;9(3):e91052.

42. Dowman R, Rissacher D, Schuckers S. EEG indices of tonic pain-related activity in the somatosensory cortices. Clin Neurophysiol 2008;119(5):1201–12.

43. Deacon RM. Measuring motor coordination in mice. J Vis Exp. 2013;75:e2609.

44. Jakkamsetti V, Scudder W, Kathote G, et al. Quantification of early learning and movement sub-structure predictive of motor performance. Sci Rep. 2021;11(1):14405.

45. Iaccarino H, Singer AC, Martorell AJ, et al. Gamma frequency entrainment attenuates amyloid load and modifies microglia. Nature. 2016;540(7632):230-235.

46. Martorell AJ, Paulson AL, Suk HJ, et al. Multi-sensory gamma stimulation ameliorates Alzheimer’s-associated pathology and improves cognition. Cell. 2019;177(2):256–271.

47. Bobola MS, Chen L, Ezeokeke CK, et al. Transcranial focused ultrasound, pulsed at 40 Hz, activates microglia acutely and reduces Aβ load chronically, as demonstrated in vivo. Brain Stimul. 2020;13(4):1014–1023.

48. Tan LL, Oswald MJ, Heinl C, et al. Gamma oscillations in somatosensory cortex recruit prefrontal and descending serotonergic pathways in aversion and nociception. Nat Commun. 2019;10(1):983.

49. Naghdi L, Ahonen H, Macario P, Bartel L. The effect of low-frequency sound stimulation on patients with fibromyalgia: a clinical study. Pain Res Manag. 2015;20(1):e21–7.

50. Kantor J, Campbell EA, Kantorova L, et al. Exploring vibroacoustic therapy in adults experiencing pain: a scoping review. BMJ Open. 2022;12(4):e046591.

51. Kim H, Park MY, Lee SD, Lee W, Chiu A, Yoo SS. Suppression of EEG visual-evoked potentials in rats through neuromodulatory focused ultrasound. Neuroreport. 2015;26(4):211–5.

52. Plaksin M, Kimmel E, Shoham S. Cell-type-selective effects of intramembrane cavitation as a unifying theoretical framework for ultrasonic neuromodulation. eNeuro. 2016;3(3):0136–15.

53. Chen G, Zhang Y, Li X, et al. Distinct inhibitory circuits orchestrate cortical beta and gamma band oscillations. Neuron 2017;96(6):1403–1418.e6.

54. He Y, Chen Y, Tian X, et al. CaMKIIα underlies spontaneous and evoked pain behaviors in Berkeley sickle cell transgenic mice. Pain. 2016;157(12):2798–2806.

55. Plone MA, Emerich DF, Lindner MD. Individual differences in the hotplate test and effects of habituation on sensitivity to morphine. Pain. 1996;66(2-3):265–70.

56. Gunn A, Bobeck EN, Weber C, Morgan MM. The influence of non-nociceptive factors on hot-plate latency in rats. J Pain. 2011;12(2):222–7.

57. Kim MG, Yoon S, Chiu CT, Shung KK. Investigation of optimized treatment conditions for acoustic-transfection technique for intracellular delivery of macromolecules. Ultrasound Med Biol. 2018;44(3):622–634.

58. Nelson TR, Fowlkes JB, Abramowicz JS, Church CC. Ultrasound biosafety considerations for the practicing sonographer and sinologist. J Ultrasound Med. 2009;28(2):139–150.

59. Darbari DS, Hampson JP, Ichesco E, et al. Frequency of hospitalizations for pain and association with altered brain network connectivity in sickle cell disease. J Pain. 2015;16(11):1077–86.

